# Increased CaMKK2 expression is an adaptive response that maintains the fitness of tumor-infiltrating natural killer cells

**DOI:** 10.1101/2022.05.18.492328

**Authors:** Patrick K. Juras, Luigi Racioppi, Debarati Mukherjee, Sandeep Artham, Xia Gao, Laura Akullian D’Agostino, Ching-Yi Chang, Donald P. McDonnell

## Abstract

Calcium/calmodulin-dependent protein kinase kinase 2 (CaMKK2) is a key regulator of energy homeostasis in several cell types. Expression of this enzyme in tumor cells promotes proliferation and migration, and expression in tumor-associated immune cells facilitates M2 macrophage polarization and the development of myeloid-derived suppressor cells. Thus, there has been considerable interest in developing CaMKK2 inhibitors as potential cancer therapeutics. However, the roles of CaMKK2 in other cellular compartments within the tumor immune environment remain to be established, an impediment to the clinical development of these agents. We report that CaMKK2 is expressed at low basal levels in natural killer (NK) cells but is significantly upregulated in tumor-infiltrating NK cells where it suppresses apoptosis and promotes proliferation. It was further demonstrated that NK cell-intrinsic deletion of CaMKK2 increased metastatic progression across several murine models, establishing a critical role for this enzyme in NK cell tumor immunity. Interestingly, ablation of the CaMKK2 protein, but not inhibition of its kinase activity, resulted in decreased NK cell survival. These results indicate an important scaffold function for CaMKK2 in NK cells and suggest that competitive CaMKK2 inhibitors and ligand-directed degraders (LDDs) are likely to have distinct therapeutic utilities. Finally, we determined that intracellular lactic acid is a key driver of CaMKK2 expression, suggesting that upregulated expression of this enzyme is an adaptive mechanism by which tumor-infiltrating NK cells mitigate the deleterious effects of a lactate-rich tumor environment. The findings of this study should inform strategies to manipulate the CaMKK2 signaling axis as a therapeutic approach in cancer.

## INTRODUCTION

Solid tumors create unique metabolic challenges that tumor-associated cells must navigate to survive and proliferate. CaMKK2 is a calcium-activated serine/threonine kinase that regulates metabolic functions through several pathways, including AMPK and CaMKIV^1,2^. Originally studied in the context of central nervous system physiology^3,4^, CaMKK2 was subsequently found to play important regulatory roles in many tissues, most notably control of gluconeogenesis in the liver^5^, insulin signaling in the pancreas^6^, and osteoblast activity in the bone^7^. Dysregulated CaMKK2 signaling also contributes to the pathology of several diseases, including non-alcoholic fatty liver disease (NAFLD) and tumor progression in many different cancers^1,2^. Indeed, ectopic expression of CaMKK2 results in the suppression of anoikis^8^, enhanced migratory capacity^9^, and increased proliferation of cancer cells^10^.

In the immune system, CaMKK2 is highly expressed in early hematopoietic stem and progenitor cells, where it promotes quiescence and suppresses differentiation^11,12^. CaMKK2 expression in granulocyte-monocyte progenitor cells promotes monocyte lineage commitment, and its expression is attenuated as granulocytes mature^11,13^. CaMKK2 remains highly expressed in monocytes and monocyte-derived macrophages, and deletion of this kinase attenuates macrophage-mediated inflammatory responses^14^. Recently, we showed that pharmacologic inhibition of CaMKK2 prevents the accumulation of M2-like tumor-associated macrophages (TAMs)^15^ while decreasing the number of myeloid-derived suppressor cells (MDSC) in the tumor microenvironment^16^. Thus, CaMKK2 contributes to tumor biology through its ability to regulate cancer cell-intrinsic biology and through its actions in specific myeloid populations^8-10,15-17^. However, the role of CaMKK2 in other tumor-infiltrating immune populations has yet to be studied, limiting our ability to develop a complete model of how CaMKK2 influences tumor pathobiology.

Given its role in promoting tumor progression, there is considerable interest in developing CaMKK2 inhibitors as cancer therapeutics. The most widely used CaMKK2 antagonist, the competitive inhibitor STO-609, has shown efficacy in mouse models of breast cancer and NAFLD^15,18^. However, the poor pharmaceutical properties of this drug coupled with its established off-target activities are an impediment to its clinical development^19^. Several new inhibitors are emerging with improved properties, and these new drugs are divided into two functional categories: ATP-competitive inhibitors of kinase function, and ligand-directed degraders (LDDs). LDDs are a new class of bi-specific molecules that feature a target-binding ligand paired with a ubiquitin ligase recruitment domain that promotes proteasomal degradation of the target protein. LDDs may offer advantages over inhibitors, including activity against inhibitor-resistant cancers and enhanced target selectivity^20,21^, though it remains to be determined whether CaMKK2-specific LDDs and competitive inhibitors are functionally equivalent. As these compounds are currently under investigation as cancer therapeutics, it has become important to understand the effects of CaMKK2 modulation on immune cell function.

## METHODS AND MATERIALS

### *In vitro* NK cell cultures

WT BL/6 mouse spleens were crushed through a 40 μm cell strainer into PBS+ 2% FBS, and the EasySep Mouse NK Cell Isolation Kit (StemCell Technologies, cat no. 19855) was used to isolate NK cells through negative selection. Unless otherwise stated, cells were cultured in a 50/50 mixture of EO771 TCM and normal NK cell growth media with IL-2 and IL-15 stimulation (Supplementary Table 1). The CaMKK2 inhibitors CC8977, GSKi^15^, and YL-36 (Sigma Aldrich, cat no. SML2834) or the LDDs CC3240 and CC3756 were added to the NK cell cultures immediately after plating.

### Generating TCM

One million EO771 or A7C11 tumor cells were plated on a 10-cm cell culture dish in their standard media and incubated at 37°C. At exactly 72 hours, the growth media was recovered from each plate, centrifuged at 2000 x g for 5 minutes, and the supernatant frozen at -80°C.

### CFSE proliferation assay

NK cells were treated with 5 μM CellTrace CFSE (Invitrogen, cat no. C34554A) in RMPI media+ 5% FBS for 5 minutes, washed twice with the same media, and cultured for 72 hours. NK cells were gated as live CD45+/CD3-/NK1.1+/CD49b+ cells.

### *In vitro* cytotoxicity assay

25,000 CFSE-stained YAC-1 cells were mixed with live NK cells at a ratio of 1:25, 1:5, 1:1, and 5:1 (NK:YAC-1) in NK cell media on a 96-well plate. The cells were co-incubated at 37°C for 4 hours and then stained with Live/dead stain and anti-CD45-BV605 for flow cytometry analysis. YAC-1 cells were identified as CD45-/CFSE+, and their viability was recorded.^22^

### *In vitro* migration assay

500,000 live NK cells were suspended in 100 μL of growth media and added to the upper insert of a transwell migration chamber with 5 μm pores (Corning, cat no. 3421). 600 μL of normal growth media containing 0 or 250 ng/mL CCL4 (BioLegend, cat no. 554602) was added to the lower chamber, and the migration plate was incubated at 37°C for 3 hours. The total number of cells in the lower chamber was measured via flow cytometry.

### Mouse strains and cell lines

The [Tg(CaMKK2-eGFP)BL/6] reporter mouse line originates from the Mutant Mouse Regional Resource Center^23^. This strain features the sequence of enhanced green fluorescent protein (eGFP), followed by a polyadenylation sequence, inserted into the mouse genomic bacterial artificial chromosome RP23-31J24 at the initiation codon of the CaMKK2 gene.

The CaMKK2-/-(KO) mouse strain and the CaMKK2^fl/fl^ mouse strain were originally generated in the lab of Anthony Means (Baylor College of Medicine)^4^. To generate NKp46-cre CaMKK2^fl/fl^ mice, the CaMKK2^fl/fl^ mice were bred with NKp46-iCre mice acquired from the lab of Joseph Sun (Memorial Sloan Kettering Cancer Center) and originally created by Eric Vivier (Centre d’Immunologie de Marseille-Luminy). To generate tamoxifen-inducible CaMKK2 KO mice, the CaMKK2^fl/fl^ mice were bred with UBC-Cre-ERT2 mice acquired from Jackson Labs (strain #007001). The ERT2+/-CaMKK2^fl/fl^ mice were dosed with 2 mg tamoxifen citrate (Sigma-Aldrich, cat no. T9262) in corn oil via oral gavage daily for one week, and splenic NK cells were harvested 5 days after dosing ended.

For additional mouse strains and cell lines see Supplementary Tables 2-3.

### Cell line modification with luciferase

HEK293T cells were grown to 40% confluency in a 10-cm plate, and the luciferase transfection mixture (Supplementary Table 4) was added dropwise after a 30 minute incubation period. 24 hours later, the media was aspirated and replaced with DMEM media containing 30% FBS. 24 hours later, the lentivirus-containing media was recovered, passed through a 45 μm filter, diluted 1:1 in DMEM, and added to B16-F10 or EO771 cell cultures. 24 hours later, the virus-containing media was aspirated, and a second round of diluted virus-containing media was added. 24 hours later, the virus-containing media was replaced with normal growth media, and transfected cells were selected using 200-1000 μg/mL neomycin (Geniticin brand, Gibco, cat no. 3480).

### Metastasis studies

Splenic NK cells were isolated from 10-12 week WT and CaMKK2 KO female mice and cultured for 24 hours in growth media containing 100 ng/mL murine IL15. 6-7 week female NSG mice were injected with 250,000 live WT or KO NK cells via the lateral tail vein in 100 μL HBSS. When using the B16-F10 model, 500,000 tumor cells were injected concurrently with the NK cells. 200 units of murine IL2 in 100 μL PBS was injected into the peritoneal cavity immediately after adoptive transfer of the NK cells and every third day thereafter. When using the β2m-/-EO771 model, 300,000 tumor cells were injected into the lateral tail vein 48 hours prior to the NK cells. Mice were regularly anesthetized with isoflurane and intravenously administered 100 μL luciferin (Regis Technologies, cat no. 1-360243-200) at the standard concentration. Bioluminescence was measured using an IVIS Lumina XR optical imaging system with an exposure time of 2-3 minutes. Lungs were extracted post-mortem, mechanically pulverized in 5 mL DMEM+5% FBS, and enzymatically digested with 1 mg/mL Collagenase A (Roche, cat no. 10103586001) and 200 units/mL DNAse I (Sigma, D5025-150KU) for 30 minutes in a 37°C shaker. Cells were stained for flow cytometry analysis.

### Primary tumor studies

11-12 week-old CaMKK2-eGFP reporter mice were injected subcutaneously with 200,000 EO771 tumor cells in 100 μL HBSS. Upon reaching a volume of 1 cm^3^, tumors and spleens were recovered and prepared for flow cytometry in the manner described above.

### Flow cytometry

Data was collected using the LSRFortessa X-20 (BD Biosciences) and analyzed using FlowJo software version 10.7.1. The Accuri C6 flow cytometer (BD Biosciences) was used for the *in vitro* migration and cytotoxic assays. For antibodies, see Supplementary Table 5.

### Western immunoblotting

Images were captured using the LI-COR Odyssey Classic scanner and processed using LI-COR Image Studio software, version 4.0. For antibodies, see Supplementary Table 6.

### Statistical analysis

Standard error of the mean is shown based on 2-3 biological replicates per experiment. Excel was used for statistical analysis. P-value was determined using an unpaired Student’s t-test with a significance threshold of P<0.05 (*P<0.05, **P<0.01, ***P<0.005).

## Data availability statement

All data generated in this study is available upon request from the corresponding author.

## Ethics statement

All animal work was approved by the Duke Institutional Animal Care and Use Committee and was conducted in a manner that minimizes pain and distress in the animal subjects.

## RESULTS

### CaMKK2 is upregulated in tumor-infiltrating mouse NK cells

As a first step in this study, we screened for CaMKK2 expression across several immune cell types under normal physiological conditions in mice. For these initial experiments we used a [Tg(CaMKK2-eGFP)BL/6] transgenic reporter mouse line expressing eGFP under the control of the CaMKK2 promoter^23^. Eight tissue types were harvested from the CaMKK2 reporter mice and subjected to flow cytometry analysis, using a previously described gating strategy to identify ten distinct immune populations **(Fig. 1A-B, S1A-B)**^15,24^. The results from this experiment confirmed that the eGFP reporter is highly active in myeloid cells, especially macrophages^14,16^. In contrast, eGFP expression was not observed in B cells or any T-cell subset **(Fig. 1A, S1A)**. Surprisingly, eGFP expression was also detected in a subset of mature NK cells, which expressed the typical myeloid marker CD11b **(Fig. 1A**)^25^. This eGFP-expressing subpopulation of CD11b^+^ NK cells was identified in blood, bone marrow, and inguinal lymph nodes, among other tissues **(Fig. 1B)**.

**Figure 1:**
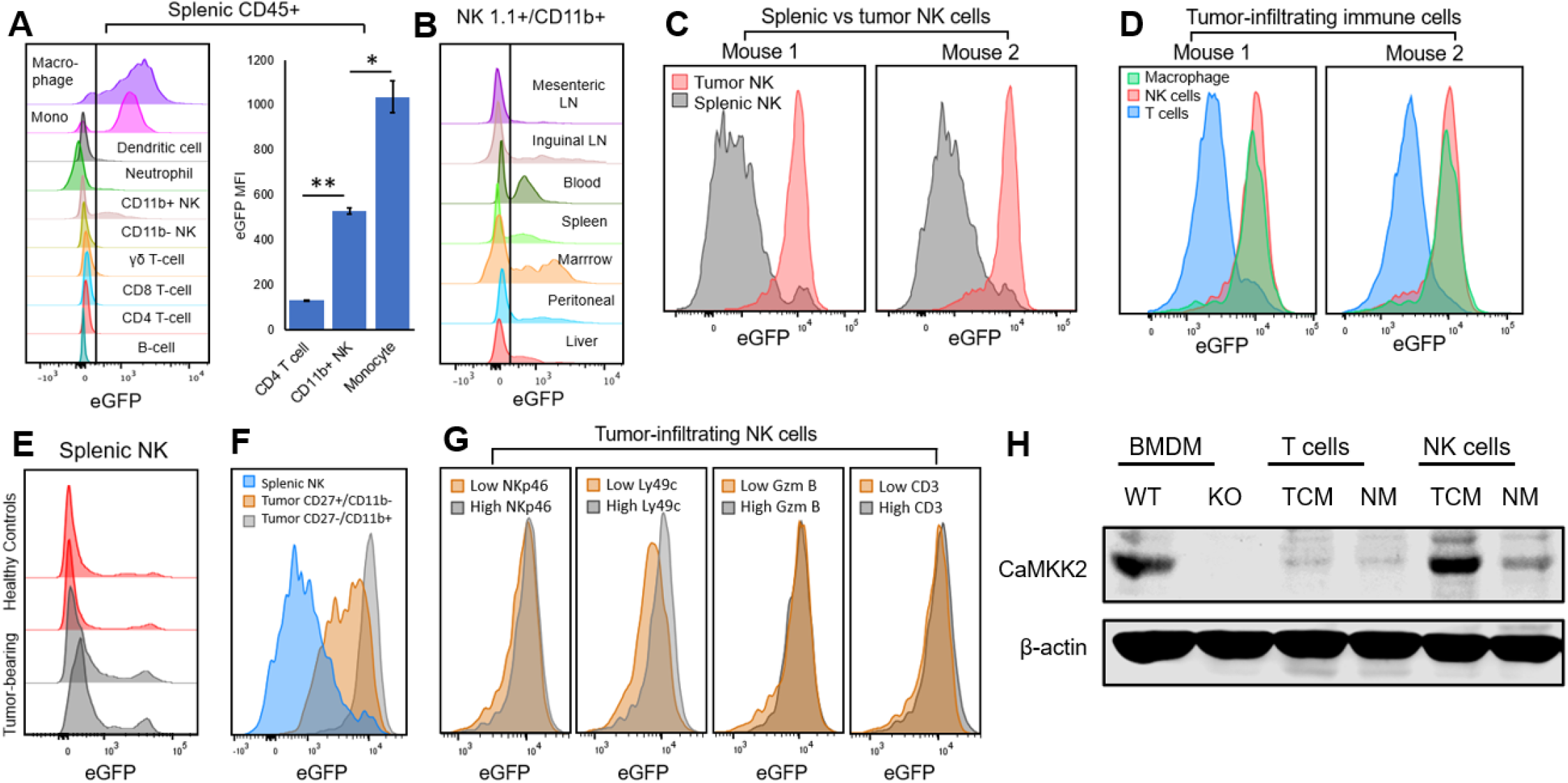
CaMKK2 is expressed in tumor-infiltrating natural killer cells. (A) Splenic immune cells from healthy CaMKK2-eGFP reporter mice were harvested and analyzed for eGFP fluorescence using flow cytometry. Each distribution represents >500 cells; data is representative of 2 mice. Standard error is shown. (B) Mature NK cells (NK1.1+/CD11b+) were isolated from various tissues of healthy CaMKK2-eGFP reporter mice and analyzed for eGFP fluorescence using flow cytometry. Each distribution represents >1000 cells; data is representative of 2 mice. (C-D) 14-day subcutaneous EO771 tumors and spleens were harvested from two CaMKK2-eGFP reporter mice. (C) NK cells from tumors or spleen, or (D) T-cells, NK cells, and macrophages from the tumor were analyzed for eGFP fluorescence using flow cytometry. (E) Splenic NK cells were harvested from CaMKK2-eGFP reporter mice bearing 14-day subcutaneous EO771 and from healthy control reporter mice, and eGFP fluorescence was analyzed using flow cytometry. (F-G) 14-day subcutaneous EO771 tumors were harvested from CaMKK2-eGFP reporter mice, and eGFP expression was analyzed on a variety of NK cell subsets. Each distribution represents >1000 cells; data is representative of 2 mice. (H) Bone marrow-derived macrophages, splenic T-cells, and splenic NK cells harvested from WT and CaMKK2-/-BL/6 female mice were cultured in 50% EO771 tumor-conditioned media (TCM) or normal growth media (NM) for 72 hours. CaMKK2 and the β-actin control were measured by Western immunoblotting following resolution of proteins on a 7.5% acrylamide gel.

We repeated the evaluation of CaMKK2 expression in tumor-infiltrating immune cells recovered from subcutaneous EO771 mammary tumors propagated in the CaMKK2 reporter mice. Notably, NK cells recovered from these tumors showed significantly higher eGFP expression than those recovered from the spleens of the same mice **(Fig. 1C)**, with levels of eGFP expression approaching the macrophage positive control **(Fig. 1D)**. eGFP expression in the splenic NK cells from tumor-bearing mice was similar to that found in healthy mice **(Fig. 1E)**, strongly suggesting that the upregulation of CaMKK2 promoter activity is specific to the tumor environment rather than a systemic response in tumor-bearing animals. Among tumor-infiltrating NK cells, expression of eGFP was correlated with, though not specific to, the later maturation states **(Fig. 1F)**. However, there was no correlation between eGFP expression and that of functional markers NKp46, Granzyme B, or Ly49c, or with the CD3+ NKT subset **(Fig. 1G)**.

To confirm these findings, we conducted Western immunoblots to directly evaluate CaMKK2 protein expression. WT splenic NK and T-cells were cultured in EO771 tumor-conditioned media (TCM) and subsequently analyzed for CaMKK2 expression. TCM is exhausted media recovered from tumor cells grown *in vitro*, and it offers a controlled approximation of the tumor environment. Bone-marrow derived macrophages (BMDMs) were harvested from WT and CaMKK2-/-(KO) mice, differentiated *in vitro* using M-CSF, and used as a positive control on the Western blots. In this manner, we confirmed that CaMKK2 is robustly expressed in tumor-conditioned NK cells compared to the low baseline expression observed in NK cells cultured in normal media **(Fig. 1H)**. Moreover, CaMKK2 protein is undetectable in tumor-conditioned T-cells **(Fig. 1H)**. These findings indicate that CaMKK2 expression is significantly upregulated in NK cells within tumors, a phenomenon that is not generalizable among lymphocytes.

### Ablation of NK cell-intrinsic CaMKK2 enhances metastatic tumor growth in multiple tumor models

Given the clinical importance of metastatic disease and the central role of NK cells in regulating metastatic progression^26^, we assessed the relevance of NK cell-intrinsic CaMKK2 in animal models of metastasis. To limit CaMKK2 ablation to the NK cell compartment, our initial experiments utilized NSG mice reconstituted with splenic NK cells from WT or CaMKK2 KO mice. At the time of transfusion, the WT and KO NK cells were qualitatively similar as assessed by the expression of activation and maturation markers **(Fig. S2C)**. To increase tumor cell sensitivity to NK cell killing, the EO771 tumor line was engineered to eliminate the expression of major histocompatibility complex class I (MHC I) using a CRISPR-based approach to delete both alleles of β2m **(Fig. S2A)**. The EO771 cells were also modified to express luciferase, enabling systemic monitoring of tumor burden using a luciferin-based *in vivo* imaging system (IVIS). To simulate metastasis, the modified EO771 cells were injected via tail vein. Within two weeks of tumor injection, mice reconstituted with KO NK cells exhibit a significantly larger systemic EO771 metastatic burden than mice reconstituted with WT NK cells **(Fig. 2B)**, suggesting that ablation of CaMKK2 in NK cells impairs their ability to eliminate metastatic cells.

**Figure 2:**
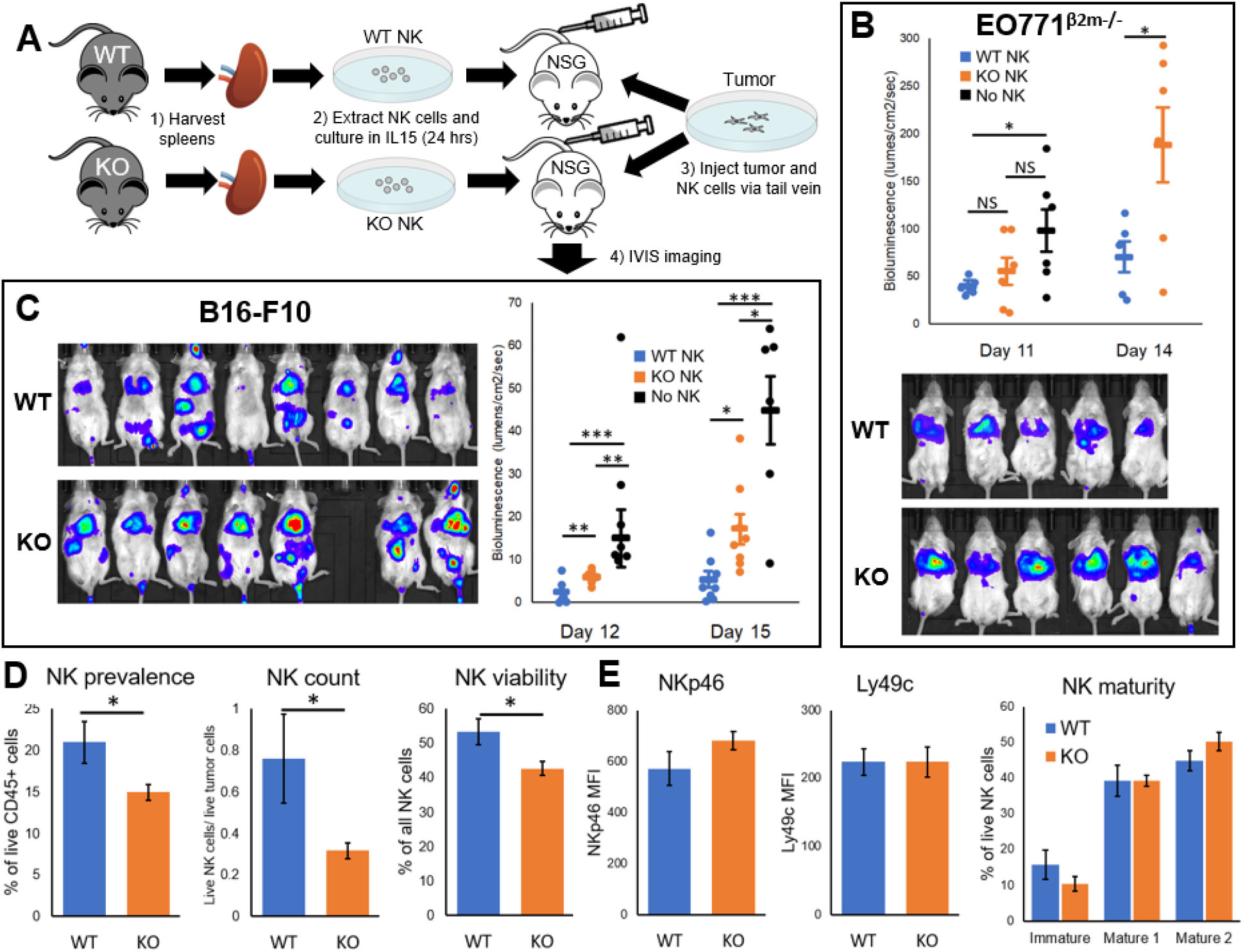
NK cells from a CaMKK2-/-background exhibit reduced activity against lung metastases. (A-C) Splenic NK cells from WT or CaMKK2-/-(KO) BL/6 females were isolated, cultured in media containing IL15 for 24 hours, and intravenously transfused into NSG mice along with luciferase-expressing B16-F10 melanoma cells or luciferase-expressing β2m-/-EO771 breast tumor cells. A control group received only tumor cells and no NK cells. (B) Whole-body metastatic burden in the EO771 study was measured using luciferin bioluminescence imaging. Results collected 14 days post-injection are shown. N=5-6, standard error shown. (C) Whole-body metastatic burden in the B16-F10 study was measured using luciferin bioluminescence imaging. Results collected 15 days post-injection are pictured. N=7-8, standard error shown. (D-E) NK cells were recovered post-mortem from the lungs of mice in the B16-F10 study and analyzed using flow cytometry. (D) Live NK cells as a portion of CD45+ cells (NK prevalence), total live NK cells as a fraction of live tumor cells (NK count), and the fraction of all NK cells that are alive (NK viability) were measured. (E) Median fluorescence intensity (MFI) of NKp46 and Ly49c on NK cells was measured. The distribution of immature NK cells (CD27+/CD11b-), Mature 1 NK cells (CD27+CD11b+), and Mature 2 NK cells (CD27-/CD11b+) was also measured. N=7-8, standard error shown.

The metastasis study was repeated using the B16-F10 melanoma model, which expresses low levels of MHC I **(Fig. S2A)**. In this model, mice reconstituted with KO NK cells also exhibited a significantly larger metastatic B16-F10 burden compared to their WT counterparts **(Fig. 2C)**. Post-mortem flow cytometry analysis of the lung tissue, which was the dominant site of metastasis, showed a significant reduction in the viability and prevalence of NK cells in the lungs of the KO group **(Fig. 2D)**. However, the WT and KO groups exhibited no differences in the expression of NK cell activation or maturation markers **(Fig. 2E)**. These results suggest that NK cell viability, but not cytotoxic function, is compromised upon CaMKK2 ablation.

The NK cells transfused in the first set of metastasis studies were harvested from animals with a germline deletion of the CaMKK2 locus, raising the possibility of confounding developmental or compensatory factors. To control for these factors, we repeated the metastasis experiments using a tamoxifen-inducible CaMKK2 KO model. This model features constitutive expression of a cre-ERT2 fusion protein that translocates to the nucleus upon tamoxifen induction, excising the floxed CaMKK2 alleles **(Fig. 3A)**. Splenic NK cells were harvested from ERT2^+^ and ERT2^-^ controls after one week of tamoxifen induction, and the NK cells were co-transfused with luciferase-expressing B16-F10 cells into NSG hosts. Knockdown of CaMKK2 in BMDMs from ERT2^+^ mice, but not in the ERT2^-^ mice, was confirmed by Western immunoblot **(Fig. S2D)**. Within two weeks of tumor injection, mice reconstituted with the ERT2^+^ NK cells showed a significantly larger systemic B16-F10 tumor burden compared to the ERT2^-^ controls **(Fig. 3B-C)**.

**Figure 3:**
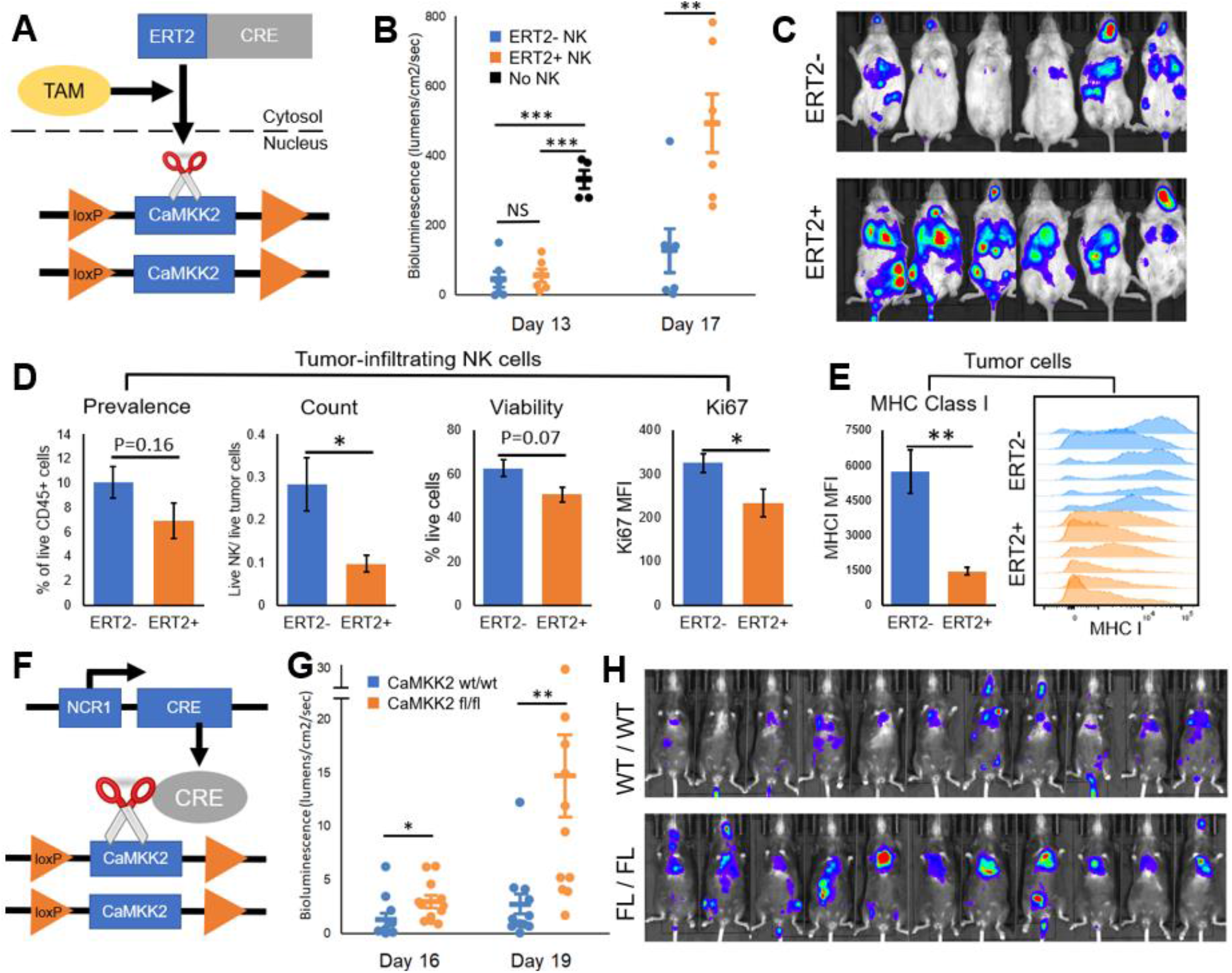
NK cell-intrinsic CaMKK2 ablation reduces NK cell survival and tumor immunoediting *in vivo*. (A-E) NK cells were harvested from the spleens of tamoxifen-treated CaMKK2^fl/fl^:ERT2+ mice or CaMKK2^fl/fl^:ERT2-mice and co-injected into the tail veins of NSG mice along with luciferase-expressing GFP+ B16-F10 tumor cells. The CaMKK2^fl/fl^:ERT2+ mice express a fusion cre-ERT2 protein that translocates to the nucleus upon tamoxifen binding, excising the floxed CaMKK2 alleles. (B-C) Full-body luciferin bioluminescence was measured over time to track metastatic progression. N=6, standard error shown. (D) NK cells were harvested from lungs 17 days post-injection. Flow cytometry was used to measure the prevalence of NK cells among live CD45+ cells (prevalence), the number of live NK cells as a proportion of live tumor cells (count), the percentage of all NK cells that are alive (viability), and Ki67 expression among NK cells (Ki67). N=6, standard error shown. (E) GFP+ tumor cells were harvested from lungs 17 days post-injection, and MHC Class I expression was analyzed via flow cytometry. N=6, standard error shown. The flow cytometry gating strategy is depicted in Figure S3B. (F-H) Mice were bred that express Cre under the NKp46 promoter and contain either CaMKK2^fl/fl^ (fl/fl) alleles or CaMKK2 wt/wt alleles, and luciferase-expressing B16-F10 tumor cells were injected via tail vein into these two groups. Full-body luciferin bioluminescence was measured over time to track metastatic progression. N=11, standard error shown.

Post-mortem flow cytometry analysis of the immune cell repertoire in the lungs showed a reduction in total live NK cells in the ERT2^+^ group compared to that observed in ERT2^-^ mice **(Fig. 3D)**, as was observed in prior studies using the germline CaMKK2 KO model. The reduction in live NK cells is likely driven by reduced proliferation, reflected in lower Ki67 expression among ERT2^+^ NK cells, and by a reduction in NK cell viability in the ERT2^+^ group **(Fig. 3D)**. We measured MHC I expression on recovered tumor cells and found a dramatic increase in the expression of this protein in the ERT2^-^ group compared to the ERT2^+^ group **(Fig. 3E)**. MHC I expression protects tumors from NK killing, so MHC I upregulation is likely the result of selective pressure exerted on the tumor cells by NK cells. Thus, persistently low MHC I expression on the tumors provides direct evidence of reduced immunoediting when NK cells lose CaMKK2 expression.

The metastasis studies conducted to this point utilized immune deficient animals in which transfused NK cells are the only functional bone marrow-derived immune population. However, many other immune populations are normally present in the tumor environment, and these populations may affect the overall phenotype or be impacted by the loss of CaMKK2 in NK cells. Thus, we developed a model in which CaMKK2 was specifically ablated in NK cells in immune competent animals **(Fig. 4F)**. Specifically, we crossed mice with floxed CaMKK2 alleles to mice expressing *cre* under the control of the NKp46 promoter, which is only active in NK cells. Experimental and control mice both express one copy of the NKp46-cre construct, but only the experimental group contain floxed CaMKK2 alleles. These mice were injected via tail vein with luciferase-expressing B16-F10 tumor cells, and within two weeks of tumor administration, the experimental group exhibited a significantly larger systemic B16-F10 tumor burden than control mice **(Fig. 3G-H)**. These studies reveal that ablation of CaMKK2 in the NK cell compartment reduces immune surveillance in immune competent animals. Extensive immune profiling of post-mortem lung tissue was performed, but we did not identify any other immunologic changes resulting from NK cell-specific CaMKK2 ablation **(Fig. S2E)**. Thus, the reduced tumor burden associated with CaMKK2 expression in NK cells is likely mediated by direct changes in the size of the NK cell population.

**Figure 4:**
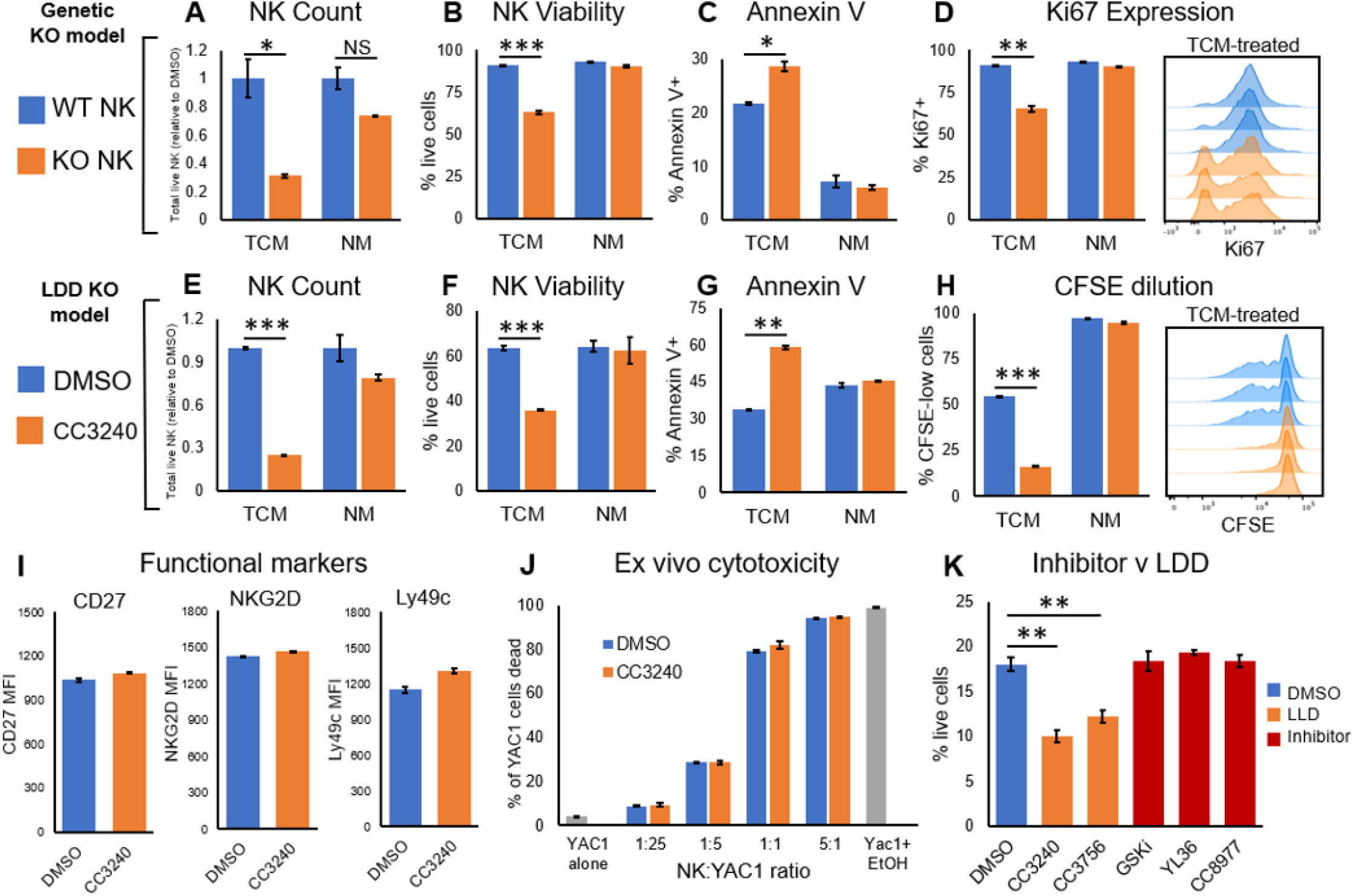
CaMKK2 ablation in tumor-conditioned NK cells reduces survival and proliferation. (A-D) Splenic NK cells were isolated from WT or CaMKK2-/-(KO) BL/6 females and cultured for 96 hours in 50% EO771 TCM or normal growth media. Flow cytometry was used to determine, (A) the total number of live NK cells as a proportion of the DMSO control, (B) the viability of the NK cells, (C) Annexin V staining on the NK cells, and (D) Ki67 expression within the NK cells. N=3, standard error shown. (E-H) Splenic NK cells were isolated from WT BL/6 females and cultured for 72 hours in 50% EO771 TCM or normal growth media containing 2.5 μM CC3240 or DMSO vehicle. Flow cytometry was used to determine, (E) the total number of live NK cells as a proportion of the DMSO control, (F) the viability of the NK cells, and (G) Annexin V staining on the NK cells. (H) A portion of NK cells were stained with CFSE dye prior to *ex vivo* culture, and CFSE dilution was measured using flow cytometry at 72 hours. N=3, standard error shown. (I) Splenic NK cells were isolated from WT BL/6 females and cultured for 72 hours in 50% EO771 TCM containing 2.5 μM CC3240 or DMSO vehicle. The median fluorescence intensity (MFI) of key functional markers was measured using flow cytometry. N=2, standard error shown. (J) Splenic NK cells were isolated from WT BL/6 females and cultured for 72 hours in 50% EO771 TCM with 2.5 μM CC3240 or DMSO vehicle. Live NK cells were co-cultured with CFSE-stained YAC-1 cells at the stated ratios for 4 hours, and YAC-1 viability was measured using flow cytometry. YAC-1 cells were cultured alone (YAC-1 alone) or with 25% ethanol (Yac1+ EtOH). N=3, standard error is shown. (K) Splenic NK cells were isolated from WT BL/6 females and cultured for 96 hours in 50% EO771 TCM containing DMSO vehicle, the LDD CC3240 or CC3756 (1 μM), or the competitive inhibitor GSKi, YL36, or CC8977 (1 μM). NK cell viability was measured using flow cytometry.

### CaMKK2 expression preserves viability and proliferative capacity of NK cells

To further elucidate the role of CaMKK2 in NK cell biology, we conducted a series of *in vitro* assays using splenic NK cells cultured in TCM. Two models of CaMKK2 ablation were employed for these *in vitro* studies: (1) NK cells harvested from WT and CaMKK2 KO mice, and (2) pharmacologic ablation of CaMKK2 in WT NK cells using a specific, high-affinity CaMKK2 LDD (CC3240). This LDD degrades about 75% of the CaMKK2 protein in primary mouse NK cells **(Fig. S3F)**.

Both the KO and CC3240-treated groups, when exposed to TCM, exhibit a significant reduction in total number of live NK cells compared to the WT and DMSO controls **(Fig. 4A, 4E)**, likely driven by a combination of increased NK cell apoptosis and reduced proliferation. When cultured in TCM, both KO and CC3240-treated NK cells showed significantly reduced viability and increased levels of Annexin V staining compared to controls **(Fig. 4C, 4G)**. Similar results were observed when TCM from A7C11 cells was used in place of the typical EO771 TCM **(Fig. S3A)**. Moreover, KO NK cells cultured in TCM exhibited reduced expression of the proliferation marker Ki67 compared to WT NK cells **(Fig. 4D)**. To directly measure proliferation, WT NK cells were stained with CFSE vital dye and cultured in TCM with CC3240 or DMSO vehicle. CFSE is diluted with each round of cell division, so reduced CFSE fluorescence serves as a proxy for proliferation. We found that the CC3240-treated NK cells were significantly less likely to undergo division compared to the DMSO-treated control **(Fig. 4H)**, further indicating that CaMKK2 expression preserves proliferative capacity. Importantly, exposure to TCM was required to elicit the noted differences in survival and proliferation **(Fig. 4A-H, S3G-I)**.

The impact of CaMKK2 on NK cell migration was also assessed. To this end, WT NK cells were cultured in TCM with CC3240 or DMSO vehicle for 72 hours, at which point equal numbers of live cells from each group were placed into the upper inserts of a migration chamber containing the CCL4 chemokine in the lower well. After three hours, it was observed that significantly fewer CC3240-treated NK cells had migrated to the bottom chamber compared to the DMSO control **(Fig. S3B)**. WT and KO NK cells express CCR5, the CCL4 receptor, at comparable levels **(Fig. S3E)**. These data, together with that from the cell survival and proliferation studies, suggest that CaMKK2 expression confers a general fitness advantage on tumor-conditioned NK cells.

The effects of CaMKK2 expression on the cytotoxic actions of NK cells were also assessed. It was determined that CaMKK2 ablation had no effect on the expression of several functional markers in TCM-treated NK cells, including NKG2D, Ly49c, and CD27 **(Fig. 4I)**. These results were not surprising given the activation marker profile on WT and KO NK cells recovered from murine lung metastases **(Fig. 2E)**. We also evaluated the direct cytotoxicity of KO or CC3240-treated NK cells. Specifically, NK cells were cultured in TCM with CC3240 or DMSO vehicle, then co-cultured with YAC-1 target cells at fixed ratios, and the survival of YAC-1 cells was measured. Neither pharmacologic ablation of CaMKK2 **(Fig. 4J)** nor genetic ablation of CaMKK2 **(Fig. S3C)** affects the cytotoxicity of NK cells against YAC-1 tumors. Together, these results suggest that CaMKK2 regulates apoptosis, proliferation, and movement of NK cells without directly affecting their cytotoxic potency.

To study the mechanism by which CaMKK2 regulates NK cell functions, we compared the effects of several functionally distinct CaMKK2-modulating compounds. CC8977, YL-36, and GSKi are high-affinity CaMKK2-specific kinase inhibitors that do not affect CaMKK2 protein levels, whereas the LDDs CC3240 and CC3756 recruit the E3 ubiquitin ligase cereblon to degrade the CaMKK2 protein, mimicking genetic KO of CaMKK2. Interestingly, only the LDDs reduce the viability of TCM-treated NK cells **(Fig. 4K)** and expression of Ki67 **(Fig. S3H)**. These results suggest a distinct scaffold function for CaMKK2 in tumor-conditioned NK cells as loss of kinase activity does not replicate the effects of protein degradation. We also treated tumor-conditioned NK cells with the most widely used competitive CaMKK2 kinase inhibitor STO609, but this drug was found to reduce NK cell survival even under normal media conditions **(Fig. S3D)**, likely reflecting its lack of target specificity at the concentrations needed for quantitative inhibition of CaMKK2^19^.

### Lactic acid exposure results in increased CaMKK2 expression in NK cells

We have shown that exposure to TCM results in a robust upregulation of CaMKK2 in NK cells, suggesting that a soluble factor(s) produced by tumor cells mediates this adaptive activity. To distinguish between protein signaling factors and small molecule factors, we repeated the *in vitro* NK cell survival assay using heat-inactivated TCM in which protein factors are denatured. NK cells cultured with heat-inactivated TCM and treated with CC3240 exhibit the same reduction in viability as NK cells cultured with normal TCM and CC3240 **(Fig. S4A)**, indicating that the factor(s) responsible for CaMKK2 upregulation is likely a non-proteinaceous small molecule. Unbiased mass spectrometry-based metabolic profiling of two TCM samples (A7C11 and EO771) was used to identify small molecule factors that may increase CaMKK2 expression^27^. In this manner, we identified twelve metabolites that were most significantly upregulated in both TCM samples compared to normal growth media **(Fig. S4B)**. The increase in lactate was particularly notable given its established role in immune cell function.

To assay the role of lactate in CaMKK2 expression, splenic NK cells from CaMKK2 reporter mice were cultured in normal growth media, EO771 TCM, or 10 mM lactic acid, a concentration found in solid tumors^28^. It was determined that eGFP expression increases significantly in the lactic acid and TCM groups, but not in the normal media group **(Fig. 5A)**. Thus, lactic acid is sufficient to induce CaMKK2 expression. Lactate could mediate this effect through several mechanisms, including GPCR engagement, acidification, or direct intracellular activity. The canonical cell-surface lactate receptor GPR81 is not expressed on any subset of NK cells **(Fig. S4D)**, but the lactate transport channel monocarboxylate transporter 1 (MCT1) is expressed in NK cells **(Fig. S4E)**. Interestingly, inhibition of MCT1 with the competitive antagonist AZD3965 prevented eGFP upregulation in CaMKK2 reporter NK cells cultured in lactic acid-rich media **(Fig. 5B)**, suggesting that lactate works intracellularly to regulate CaMKK2 expression.

**Figure 5:**
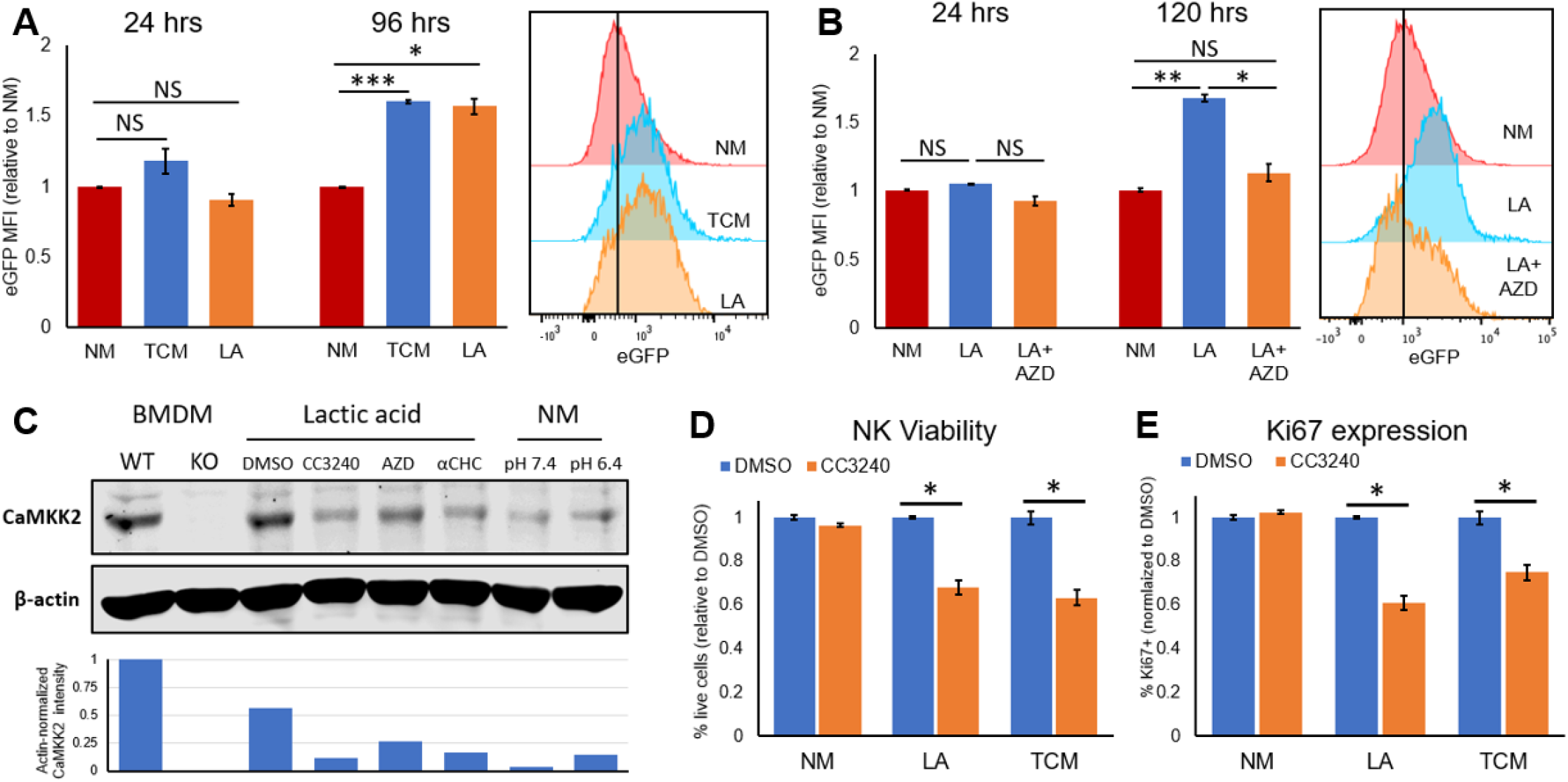
Lactic acid exposure causes CaMKK2 upregulation in NK cells. (A) Splenic NK cells isolated from CaMKK2-eGFP reporter mice were cultured for 24-96 hours in normal growth media (NM), 50% EO771 tumor-conditioned media (TCM), or normal media with 10 mM lactic acid (LA). The level of eGFP expression relative to the NM group is shown at each timepoint. The eGFP fluorescence peaks at 96 hours are shown. N=3, standard error shown. (B) Splenic NK cells from CaMKK2-eGFP reporter mice were cultured in normal media with or without 10 mM lactic acid and 500 nM AZD3965 (“AZD”) for 24-120 hours, and eGFP expression relative to the NM group is shown. The eGFP fluorescence peaks at 120 hours are shown. N=3, standard error shown. (C) Bone marrow-derived macrophages (BMDMs) were harvested from WT and CaMKK2 KO mice and differentiated in normal media with M-CSF. Splenic NK cells were harvested from WT BL/6 females and cultured in heavily buffered normal media with a fixed pH of 6.4 or 7.4 (“NM”) or in heavily buffered normal media with a fixed pH of 7.4 containing 10 mM lactic acid (“Lactic acid”). Various samples of lactic acid-cultured NK cells were simultaneously cultured with 2.5 μM CC3240, 500 nM AZD3965 (“AZD”), 5 mM αCHC (αCHC), or DMSO vehicle. CaMKK2 and the B-actin control were measured via Western immunoblotting on a 7.5% acrylamide gel (45 ug protein per lane). (D-E) Splenic NK cells from WT BL/6 females were cultured in normal media (NM), 50% EO771 TCM (TCM), or normal media with 10 mM lactic acid (LA), along with 2.5 μM CC3240 or DMSO vehicle. (E) NK cell viability and (F) expression of Ki67 are shown at 96 hours. N=3, standard error shown.

Using Western immunoblots, it was determined that lactic acid-treated NK cells robustly express CaMKK2 protein compared to the modest baseline levels in cells cultured in normal media **(Fig. 5C)**. Moreover, treating the NK cells with AZD3965 or with the pan-MCT inhibitor α-cyano-4-hydroxycinnamic acid (αCHC) significantly reduces CaMKK2 expression **(Fig. 5C)**. To control for the effects of media acidification, the NK cells used in these studies were cultured in heavily buffered media (pH of 7.4), and no upregulation of CaMKK2 was observed when NK cells were cultured in media with a fixed pH of 6.4 **(Fig. 5C)**. These data suggest that CaMKK2 upregulation is the product of direct lactic acid signaling (or metabolism) rather than an indirect effect of media acidification. The role of lactate signaling in CaMKK2 modulation was further demonstrated using survival and proliferation assays comparing TCM and lactic acid. Lactic acid reduces the viability of CC3240-treated NK cells in the same manner as TCM **(Fig. 5D)**, while also reducing expression of the Ki67 proliferation marker in CC3240-treated NK cells **(Fig. 5E)**. These data are consistent with the hypothesis that upregulation of CaMKK2 expression is a response to intra-tumoral lactic acid accumulation.

## DISCUSSION

MHC I is expressed on all nucleated cells to present intracellular antigens for immune interrogation, but many tumors downregulate expression of this ubiquitous complex to hide their mutated neo-antigens from cytotoxic T-cells^29^. Such tumors become targets for NK cells, which attack MHC I-low cells expressing markers of DNA damage common in tumors, such as MICA, MICB, and ULBP1-3^30^. NK cells also mediate antibody-dependent cellular cytotoxicity (ADCC), a process that destroys antibody-opsonized targets^31,32^. However, pro-tumor NK cell subsets have also been identified, including “decidua-like” NK cells that produce high levels of pro-angiogenic factors^33^ and “regulatory” NK cells that suppress immune activity through cytokine production^34^.

Given their multi-faceted role in tumor immunity, there has been considerable interest in manipulating NK cells for therapeutic benefit, but there are significant challenges to this approach. Transfusions of autologous cytokine-primed NK cells or highly active human NK cell lines, such as the lymphoma line NK92, have been attempted with some success^35,36^. However, immune suppression within the tumor environment is a key limiting factor^37^. Solid tumors are often hypoxic, nutrient-depleted, and awash with immunosuppressive factors such as TGFβ and activin, which disable NK cell activation and proliferation^38,39^. Long considered a waste product of anerobic metabolism, lactic acid is now appreciated as an important immunosuppressive signaling molecule in the tumor microenvironment. Lactate promotes M2 macrophage polarization through HIF1α signaling^40^, reduces MHC class II expression on dendritic cells^41^, and suppresses the activation and survival of T-cells and NK cells^42^.

Interestingly, we have determined that lactic acid facilitates the upregulation of CaMKK2 expression in NK cells, enhancing survival while facilitating proliferation, and thereby mitigating the suppressive effects of the lactate-rich tumor environment. Our data specifically support a role for intracellular lactate in the regulation of CaMKK2 as the actions of lactate (a) are blocked by inhibitors of the MCT lactate transporters, (b) are not mimicked by general acidification, and (c) do not appear to require cell surface lactate receptors. We have not yet identified the mechanisms by which intracellular lactate modulates CaMKK2 expression, although work from other groups is instructive in this regard. Lactic acid may exert direct effects on transcription by (a) functioning as a competitive inhibitor for several histone deacetylases (HDACs)^43^, (b) regulating the activity of HIF-1α^44^, or (c) directly modifying histone function through lactylation of lysine residues^45^.

While a large body of evidence indicates that NK cells play a pivotal role in the anti-tumor immune response, the function of NK cells in the primary tumor varies with tumor type^46,47^. More consistent evidence indicates a key role for NK cells in metastatic disease^26^. For this reason, we investigated the function of NK-cell intrinsic CaMKK2 in the context of tumor metastasis. Studies performed using two different tumor lines and three models of CaMKK2 ablation demonstrated that deletion of CaMKK2 in NK cells results in decreased survival and proliferation of NK cells, enabling metastatic progression. We have not probed the role of CaMKK2 in the function of human NK cells, but ongoing single cell mRNA sequencing of human breast tumors in our laboratory will likely be informative in this regard. Interestingly, the human NK lymphoma cell line NK92 expresses CaMKK2 constitutively **(Fig. S4F)**, which may reflect the selective advantage that CaMKK2 confers on the survival and proliferation of NK cells.

The contrasting effects of CaMKK2 inhibitors and LDDs on NK cell survival suggest a distinct “scaffold” function for CaMKK2. Several proteins are known to bind CaMKK2, including the eponymous calmodulin activator and the inhibitory 14-3-3γ protein^1,48^. Interestingly, CaMKK2 serves an important scaffold function in hepatocellular carcinoma, nucleating the assembly of a CaMKIV, mTOR, and S6K complex that promotes tumor proliferation^49^. This specific scaffold complex is unlikely to account for CaMKK2 activity in NK cells as it also requires the kinase function of the enzyme, but the possibility remains that the mTOR pathway may be involved given its central role in NK cell biology.

This work has significant therapeutic implications as it reveals potential liabilities associated with inhibition of MCT1 and CaMKK2, while suggesting novel strategies for enhancing NK cell immunity. By blocking the accumulation of intracellular lactate in NK cells, MCT1 inhibitors may reduce CaMKK2 upregulation, limiting the utility of this class of drugs in NK cell-sensitive tumors. In a similar manner, the detrimental effects of LDD-mediated CaMKK2 ablation in NK cells may attenuate the established benefits of CaMKK2 ablation in tumor cells and TAMs^15^. However, we did not observe any negative effects of competitive CaMKK2 inhibitors on NK cells, indicating that loss of the CaMKK2 scaffold function, but not inhibition of kinase function, is deleterious to NK cell immunity. Thus, competitive inhibitors of CaMKK2 may be clinically preferable to LDDs. CaMKK2 LDDs have yet to be adapted for *in vivo* use, but when suitable compounds become available, the efficacy of inhibitors and LDDs in NK cell-sensitive mouse tumor models will be explored. NK cells cell transfusions are being investigated as a cancer therapy, and our findings suggest that modification of these immune cells to constitutively express CaMKK2 may confer proactive resistance to the suppressive effects of the tumor environment. NK92 cells already robustly express CaMKK2, but primary human NK cells are also used for transfusions. Primary NK cells have been modified via retroviral transfection to overexpress receptors beneficial to anti-tumor activity^50^, and CaMKK2 could be artificially overexpressed using the same techniques.

## CONCLUSION

We have shown that CaMKK2 is conditionally upregulated in tumor-conditioned NK cells due to lactic acid signaling, preserving NK cell viability and proliferation in the suppressive tumor environment. Indeed, CaMKK2 expression enhances NK cell-mediated immune surveillance, slowing metastatic progression in animal models. Our work suggests that CaMKK2 mediates these effects in NK cells through a scaffold function, providing the first evidence that CaMKK2 inhibitors and LDDs may confer different benefits in cancer treatment. Overall, our work elucidates a novel adaptive mechanism by which NK cells respond to the metabolic challenges of a lactate-rich environment and provides actionable insights of therapeutic relevance.

## ACKNOWLEDGEMENTS

This work was supported by NIH grant 1F30CA261005, NCI grant K99/R00 CA237618, Department of Defense grant W81XWH-20-1-0497, and the Duke Medical Scientist Training Program. We thank the Duke Cancer Institute Flow Cytometry Core, the Duke Optical Molecular Imaging and Analysis Service Center, the Duke Cancer Center Isolation Facility, and the Duke Molecular Physiology Institute for use of their facilities and equipment. We thank Bristol-Myers Squibb for providing compounds CC3240, CC3756, and CC8977 and Dr. David Drewry at the University of North Carolina for providing YL-36.

