## Supplementary figures for "Increased CaMKK2 expression is an adaptive response that maintains the fitness of tumor-infiltrating natural killer cells"

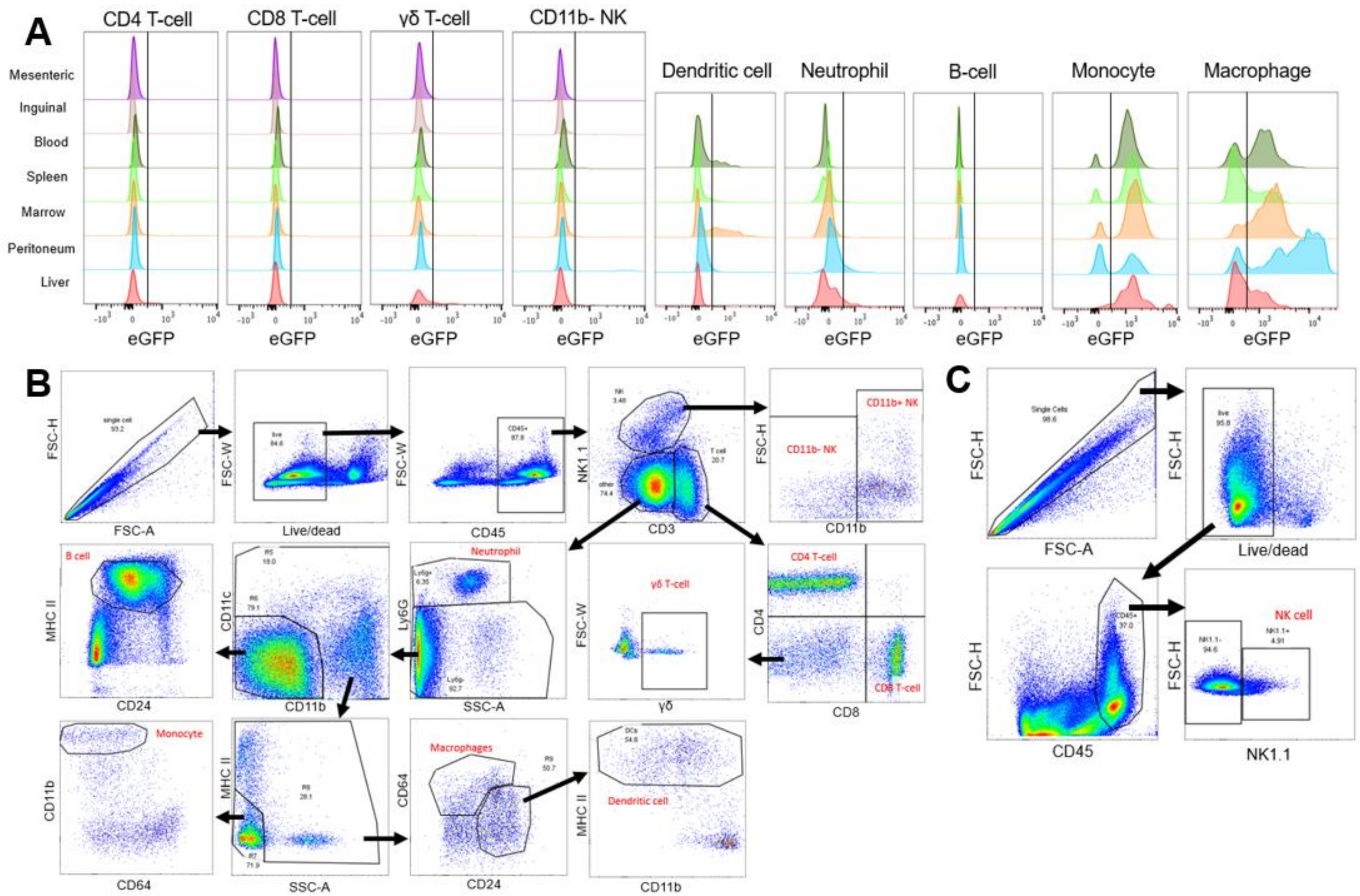


**Figure S1:** (A) Various tissues were harvested from healthy CaMKK2-eGFP reporter mice, and several immune subsets were stained for flow cytometry analysis of eGFP fluorescence. Each distribution represents >500 cells; data is representative of two mice. (B) The flow cytometry gating strategy is shown for data presented in Figure 1A-B, Figure S1A, and Figure S2E. (C) The flow cytometry gating strategy is shown for data presented in Figure 1C-G.

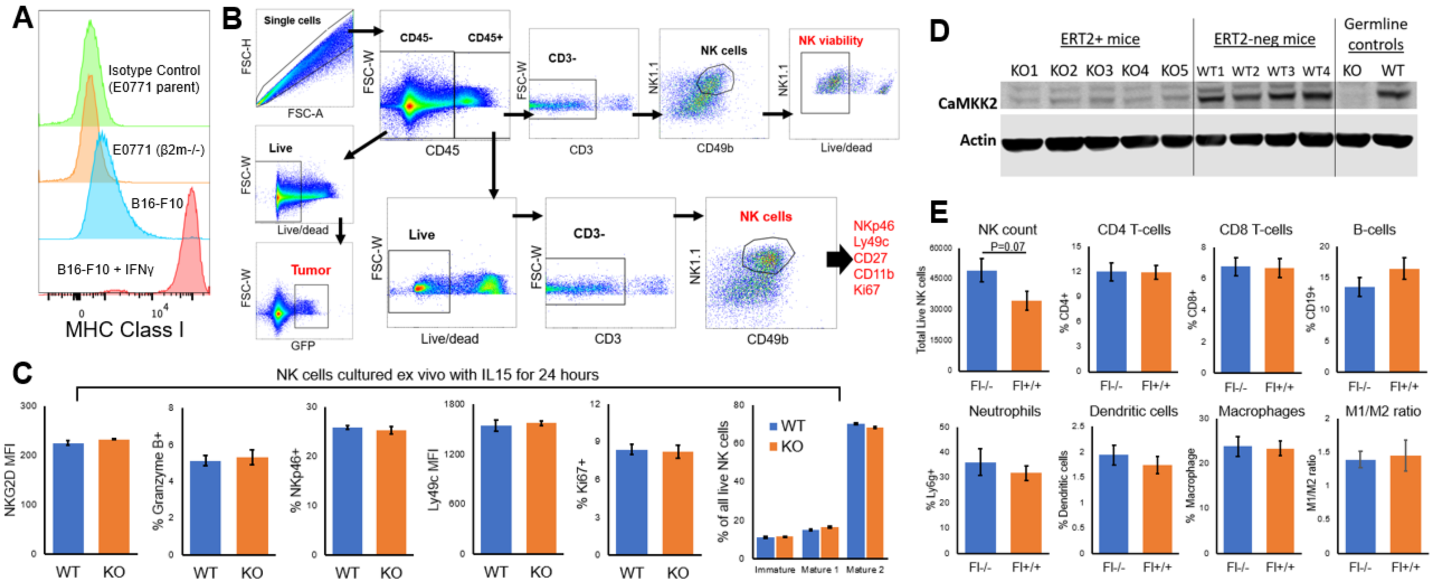


**Figure S2:** (A) β2m-/- EO771 cells, B16-F10 cells, and B16-F10 cells treated with 100 ng/mL IFNγ for 48 hours were analyzed for MHC Class I expression via flow cytometry. Fluorescence was compared to EO771 parent cells stained with an isotype control antibody. (B) The flow cytometry gating strategy is shown for data presented in Figure 2 and Figure 3D-E. (C) Splenic NK cells were harvested from WT and CaMKK2 KO BL/6 females and cultured for 24 hours in normal media containing 100 ng/mL IL15. Various functional markers were characterized via flow cytometry. The prevalence of immature NK cells (CD27+/CD11b-), mature 1 NK cells (CD27+/CD11b+), and mature 2 NK cells (CD27-/CD11b+) was also measured. N=4, standard error shown. (D) After one week of tamoxifen dosing, BMDMs were recovered from the ERT2+ mice and ERT2-negative controls to confirm CaMKK2 ablation via Western immunoblot on a 7.5% acrylamide gel. (E) In the study described in Figure 3F-H, NK cells were harvested from lungs 19 days post-injection. Flow cytometry was used to measure the absolute number of live NK cells and the prevalence of various other immune populations within the lungs. The flow cytometry gating strategy is shown in Figure S1B. N=11, standard error is shown.

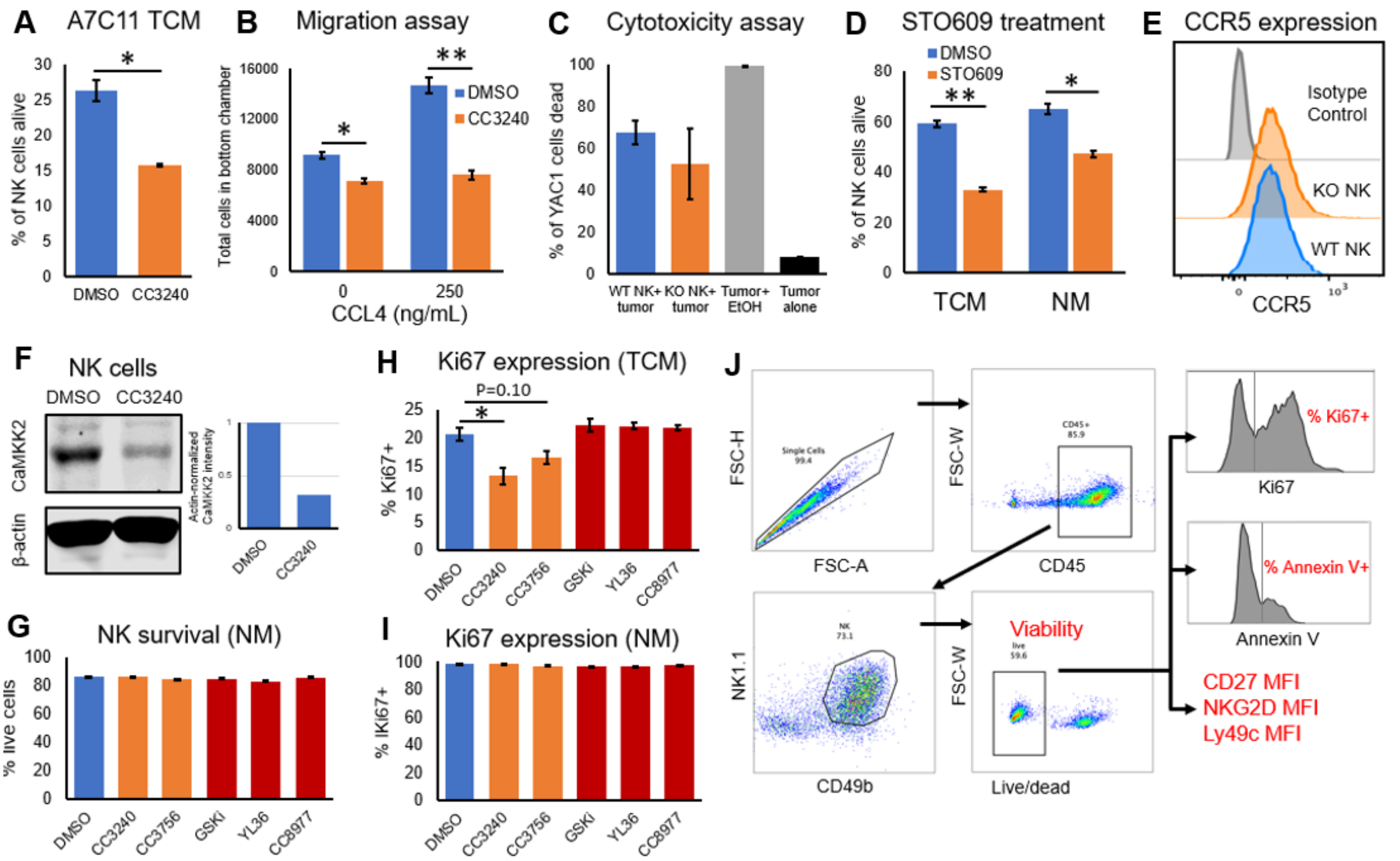


**Figure S3:** (A) Splenic NK cells were isolated from WT BL/6 females and cultured for 72 hours in 50% A7C11 TCM containing 2.5 μM CC3240 or DMSO control. NK cell viability was measured via flow cytometry. N=2, standard error shown. (B) Splenic NK cells were isolated from WT BL/6 females and cultured for 72 hours in 50% EO771 TCM containing 2.5 μM CC3240 or DMSO vehicle. 500k live NK cells were transferred to the upper chambers of a migration plate, separate from CCL4-containing media by a barrier with 5 μm pores. After 3 hours, cell number in the bottom chamber was measured via flow cytometry. N=3, standard error shown. (C) Splenic NK cells were harvested from WT or CaMKK2 KO mice and cultured in 50% EO771 media for 96 hours. Live NK cells were co-cultured with CFSE-stained YAC-1 cells at a 1:5 ratio for 4 hours, and YAC-1 viability was measured via flow cytometry. N=3, standard error is shown. (D) Splenic NK cells were isolated from WT BL/6 females and cultured for 72 hours in 50% EO771 TCM or normal growth media containing 10 μM STO609 or DMSO vehicle. NK cell viability was measured via flow cytometry. N=2, standard error shown. (E) Splenic NK cells were harvested from WT or CaMKK2 KO BL/6 females and cultured for 96 hours in normal growth media containing 100 ng/mL IL2 and IL15. CCR5 expression was measured via flow cytometry and compared to an isotype control antibody. (F) Splenic NK cells were harvested from WT BL/6 females and cultured for 72 hours in 50% EO771 tumor-conditioned media containing 2.5 μM CC3240 or DMSO vehicle. CaMKK2 and the β-actin control were measured via Western immunoblotting on a 7.5% acrylamide gel, and the actin-adjusted CaMKK2 band intensities were quantified. (G-I) Splenic NK cells were isolated from WT BL/6 females and cultured for 96 hours in 50% EO771 TCM or normal media (NM) containing either DMSO vehicle, the LDD CC3240 or CC3756 (1 μM), or the competitive inhibitor GSKi, YL-36, or CC8977 (1 μM). NK cell viability and Ki67 expression was measured via flow cytometry. (J) The flow cytometry gating strategy is shown for all data presented in Figure 4 and Figure S3.

### Figure S4

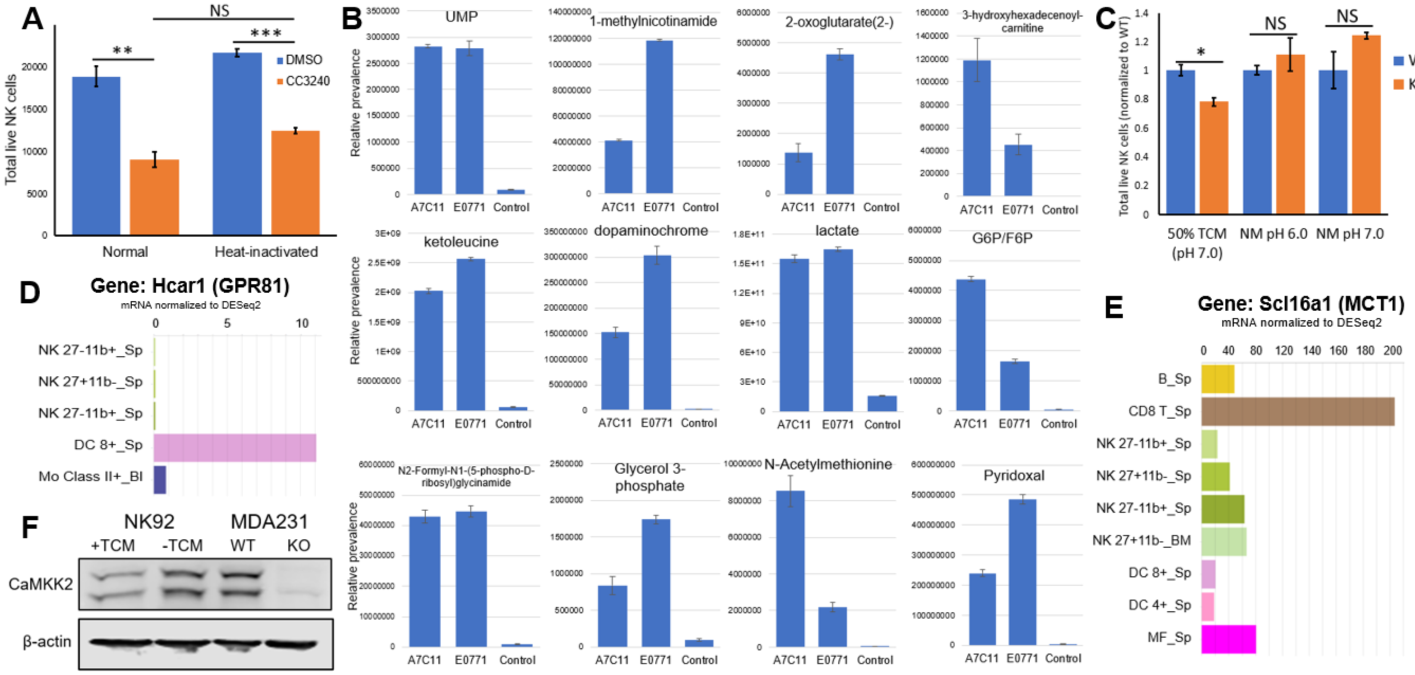


**Figure S4:** (A) Splenic NK cells from WT BL/6 females were cultured in 50% E0771 TCM (Normal TCM) or 50% E0771 TCM that had been incubated at 56 degrees for 30 minutes (Heat-inactivated TCM). Cells were treated with either 2.5 μM CC3240 or DMSO vehicle for 72 hours, and total number of live NK cells was measured via flow cytometry. N=2, standard error shown. (B) A7C11 TCM, E0771 TCM, and normal growth media (control) were analyzed via mass spectrometry for 217 common metabolites. Eight metabolites, shown in the panel, were significantly elevated in both A7C11 TCM and E0771 TCM compared to the control growth media. N=3, standard error shown. (C) Splenic NK cells were harvested from WT and CaMKK2 KO BL/6 females and cultured for 96 hours in 50% EO771 TCM, heavily buffered normal media with a pH of 6.0, or heavily buffered normal media with a pH of 7.0. The total number of live NK cells, normalized to the WT level, was measured via flow cytometry. N=2, standard error shown. (D-E) The Immunological Genome Project (Immgen) database was used to assess levels of (D) Hcar1 mRNA (codes for GPR81), or (E) Scl16a1 mRNA (codes for MCT1). (F) NK92 cells cultured in normal growth media or 50% EO771 TCM, and WT or CaMKK2-/- (KO) MDA231 cells were run on a 7.5% acrylamide cell. CaMKK2 and β-actin were detected via Western immunoblotting.

| Table 1. NK cell culture conditions | |
|---|---|
| **Reagent** | **Supplier** |
| RMPI 1640 base media | Gibco |
| 10% fetal Bovine serum (FBS) | Gibco |
| 1 mM sodium pyruvate (NaPy) | Gibco |
| 0.1 mM non-essential amino acides (NEAA) | Gibco |
| 10 mM HEPES | Gibco |
| 2 mM L-glutamine | Gibco |
| 55 µM 2-mercaptoethanol | Gibco |
| 100 units/mL Penicillin+ 100 µg/mL Streptomycin | Gibco |
| 100 ng/mL murine IL2 | PeproTech |
| 100 ng/mL murine IL15 | PeproTech |
| **If strict pH stabilization is required add:** | |
| 25 mM HEPES | Gibco |
| 25 mM PIPES | Gibco |

| Table 2. Cell line origins | |
|---|---|
| **Cell line** | **Source** |
| EO771 | Mark Dewhirst (Duke University) |
| A7C11 | Jose Conejo-Garcia (Moffitt Cancer Center) |
| B16/F10 | Duke Cell Culture Facility |
| YAC-1 | Duke Cell Culture Facility |
| HEK293T | ATCC |

| Table 3. Mouse Strain Origins | |
|---|---|
| **Mouse strain** | **Source** |
| [Tg(CaMKK2-eGFP)BL/6] | Mutant Mouse Regional Resource Center (NIH) |
| CaMKK2-/- (KO) | Anthony Means (Baylor College of Medicine) |
| CaMKK2 $^{fl/fl}$ | Anthony Means (Baylor College of Medicine) |
| NKp46-iCre | Eric Vivier (Centre d'Immunologie de Marseille-Luminy) |
| NSG mice | Jackson Labs (strain # 005557) |
| UBC-Cre-ERT2 | Jackson Labs (strain #007001) |

| Table 4. Tranfection reagents | | | |
|---|---|---|---|
| **Reagent** | **Quanitity** | **Supplier** | **Cat no.** |
| Opti-Mem | 263 µL | Gibco | 51985091 |
| FuGENE-6 | 17 µL | Promega | E2692 |
| VSVG envelope vector[1] | 280 ng | Addgene | plasmid #8454 |
| PsPAX2 Lentivirus packaging vector[2] | 2.8 µg | Addgene | plasmid #12260 |
| Lenti-luciferase P2A-Neo[3] | 2.8 µg | Addgene | plasmid #105621 |

1. Stewart SA, et al. RNA 2003 Apr;9(4):493-501.
2. psPAX2 was gifted to Addgene by Didier Trono
3. Xu Y, et al. Cancer Cell. 2018 Jan 8;33(1):13-28.e8. doi: 10.1016/j.ccell.2017.12.002.

| Table 5. Flow cytometry antibodies | | | |
|---|---|---|---|
| **Target** | **Fluorochrome** | **Supplier** | **Cat no.** |
| CD16/32 | None (blocking) | Invitrogen | 14-0161-85 |
| CD16.2 | None (blocking) | BioLegend | 149502 |
| AnnexinV | AF488 | Invitrogen | A13201 |
| CD45 | BV605 | Invitrogen | 103139 |
| CD49b | BV650 | BD Biosciences | 740496 |
| NK1.1 | AF700 | Invitrogen | 56-5941-82 |
| CD3 | PerCP/Cy5.5 | BD Pharmigen | 560527 |
| NKp46 | BV711 | BioLegend | 137621 |
| NKG2D | FITC | BioLegend | 115711 |
| Ly49c | PE/Cy7 | BioLegend | 108210 |
| CD11b | PE | BioLegend | 101208 |
| CD27 | BV786 | BioLegend | 124241 |
| CCR5 | PE | Invitrogen | 12-1951-81 |
| CD24 | BUV496 | BD Horizon | 612953 |
| CD64 | PerCP/Cy5.5 | BioLegend | 139308 |
| IA/IE | BV711 | BD Horizon | 563414 |
| CD11c | APC/Cy7 | BD Pharmigen | 561241 |
| CD11b | AF700 | BioLegend | 101222 |
| Ly6G | BV785 | BioLegend | 127645 |
| Ly6C | BV711 | BioLegend | 128037 |
| Granzyme B | AF647 | BioLegend | 515406 |
| Ki67 | BUV395 | BD Horizon | 564071 |
| CD206 | FITC | BioLegend | 141710 |
| LIVE/DEAD Fixable Violet Dead Cell Stain | BV421 | Invitrogen | L34964 |

| Table 6. Western antibodies | | | | |
|---|---|---|---|---|
| **Target** | **Clone** | **Supplier** | **Cat no.** | **Dilution** |
| CaM Kinase Kinase | 6/CaM Kinase | BD Transduction Labs | 610544 | 1:1,000 |
| β-actin | AC-15 | Sigma-Aldrich | A1978 | 1:10,000 |
